# Anthropogenic factors explain wing colour variation in *Colias* (Lepidoptera: Pieridae) butterflies across the Toronto urbanization gradient

**DOI:** 10.1101/2025.11.06.686962

**Authors:** Amanda Sabatino, Joshua P. Jahner, S. Eryn McFarlane

## Abstract

Hybridization, or interbreeding between two previously diverged populations, is increasing due to human influences on the environment. Rates of hybridization might be increasing particularly quickly among urban species, especially those that are known to be sensitive to environmental disturbances. Butterflies are one such taxa, as some species are commonly found in urban areas, despite being sensitive indicators. For example, *Colias philodice* and *C. eurytheme* (Lepidoptera: Pieridae) hybridize when their ranges are in contact or overlap and are commonly found in large cities throughout North America. However, it is an open question how variation in urbanization might affect variation in hybridization. Using wild *Colias* across a gradient of urbanization in Toronto, Ontario, we tested how the variation in *Colias* colour, as an indicator of hybridization, relates to human disturbance and urbanization. We created an objective numeric classification for both *C. eurytheme* and *C. philodice*, against which we categorized *Colias* samples. We found that some of the urbanization metrics, including distance to road, Julian date, and number of pedestrians affect the variation in *Colias* colour, suggesting that urbanization affects *Colias* colouration, and potentially rates of hybridization, in Toronto. This means that human disturbance of the environment could be affecting *Colias* rates of hybridization.

## Introduction

Hybridization is the interbreeding that occurs between two evolutionary lineages, with individuals being distinguishable based on one or more characteristics (Harrison, 1990; Harrison and Larson, 2014). Approximately 10% of animal species hybridize (Mallet et al., 2007; Aubert et al., 1997), which can result in introgression, or the integration of alleles from the gene pool of one species into another (Harrison and Larson, 2014; Anderson and Hubricht, 1938). While introgression can both increase phenotypic variation within backcrossed populations (Rieseberg et al., 1999; Grant and Grant, 2008), it can also decrease biodiversity, and thus is considered a conservation concern (Rhymer and Simberloff, 1996; Allendorf et al., 2001). Finally, human-induced environmental changes may alter both the rates, patterns, and outcomes of hybridization.

Habitat fragmentation and other human-induced changes to the environment, such as climate change, can directly affect hybridization dynamics (Ålund et al., 2023; Todesco et al., 2016; Larson et al., 2019). Alterations to the environment influenced by human-induced changes may enhance the fitness of hybrids and increase niches that enhance hybrid fitness (Todesco et al., 2016), although examples of specific human influences on hybridization rates are fairly rare. One exception is that (Grabenstein et al., 2023) found that Black-capped (*Poecile atricapillus*) and Mountain chickadees (*P. gambeli*) hybridize at increased rates depending on levels of human disturbance. Similarly, historical rates of stocking and other environmental variables have been found to affect rates of hybridization between Yellowstone cutthroat (*Oncorhynchus clarkii bouvieri*) and rainbow trout (*O. mykiss*) (Mandeville et al., 2019). Given the rapid rate of urbanization (Sudha et al., 2012), as a result of urban densification and its effect on the environment, and how urbanization produces fragmented habitats (Sudha et al., 2012), there is good reason to suspect that urbanization could generally affect hybridization rates.

Urbanization increases pressures on ecosystems (Grimm et al., 2008), and thus can affect both rates of hybridization and selection on hybrid individuals. Organisms in cities are faced with urbanization at varying degrees, with a wide range of outcomes (Concepción et al., 2017). In ecosystems that are highly disturbed, like cities, human disturbance could change the fitness landscape to one that favours hybrids (McFarlane and Mandeville, 2023; Sartor et al., 2021). The high spatial heterogeneity of urban environments helps promote some species, allowing them to be found in a greater abundance within urban habitats (Savard et al., 2000; McKinney, 2008). Human disruptions within areas of high urbanization, result in spatial isolation, promote genetic diversity, and also can promote hybridization (Ålund et al., 2023). Human changes to the environment can advance the establishment of new ecological niches or the removal of niches, causing varying selection (Ålund et al., 2023). For example, two species of red wood ants *Formica polyctena* and *F. rufa*, which typically do not interbreed with one another, were found to have an increased presence of intermediate individuals in areas that were highly disturbed and fragmented (Seifert et al., 2010). Recognizing the potential effects of human-induced landscape changes, particularly urbanization, on ecosystems and altered niches is important in understanding varying hybridization rates.

Butterflies are excellent models in urban settings as they are sensitive to urbanization (Forister et al., 2023), but are still persistent in city settings. Specifically, butterflies are suitable indicators of indirect measures of environmental variation: as they are highly responsive to local climate and light levels, and their patterns are important in studying habitat shifts (Snell-Rood et al., 2014; Blair and Launer, 1997). The intensity and effect that urban environments have on butterflies varies between species (Hardy and Dennis, 1999). For example, generalist species tend to be more common in areas with greater urban pressures (Bergerot et al., 2011). The size of parks can even affect butterfly diversity in cities, with larger parks containing more species (Sing et al., 2016). Taken together, butterflies are expected to be responsive to urbanization, and make a good model for understanding how the various, related human pressures in a city can affect rates of hybridization. How *Colias* respond to urbanization is an open question. *C. eurytheme* and *C. philodice* hybridize where their ranges are in contact or overlap, producing F1 hybrids consisting of intermediate characteristics (Hovanitz, 1949). Furthermore, there is the production of backcross offspring and F2 hybrids (Hovanitz, 1949). Hybridization that occurs between the two species does not eliminate either, rather *C. eurytheme* and *C. philodice* continue to persist regardless of gene exchange (Hovanitz, 1949; Dwyer et al., 2015). The relative abundance of hybrids between *C. eurytheme* and *C. eriphyle* is influenced by warmer temperatures due to varying seasonal weather variables affecting abundance (Jahner et al., 2012). Thus, with a rise in temperatures in urban environments, we might expect increased hybridization between *Colias*. Additionally, recent genetic work has shown that wing colouration is controlled by two major-effect loci and one minor-effect locus which explain over 70% of the heritable variation in colour, suggesting an oligogenic architecture (Hanly et al., 2023). Hybridization between *C. eurytheme* and *C. philodice* can therefore be assessed through a phenotypic approach, as introgression occurs through visible traits, with hybridization being assumed as the presence of intermediates (Jahner et al., 2012; Hovanitz, 1949). Thus, wing colouration can be used as a proxy of hybridization between *C. eurytheme* and *C. philodice* (Hanly et al., 2023).

As far as we are aware, it is unknown how urbanization affects hybridization in *Colias*. Using a hypothesis-generating research approach, our study incorporated a variety of data including temperature, pedestrian density, date, community density, the distance to the nearest tree, and the distance to the nearest road as measures of urbanization to determine relationships among urbanization and *Colias* phenotype. If urbanization affects hybridization, then we predicted that the proportions of intermediate phenotypes would increase along an urbanization gradient, reflecting the elevated rates of hybridization of *C. philodice* and *C. eurytheme*.

## Methods

### Creating an objective classification

Hybridization between species can be detected through genomic tools and phenotypic measures, allowing for ancestry to be assessed without the need to directly observe mating (Grabenstein and Taylor, 2018). In the study conducted by Hovanitz (1949), individuals were classified based on their orange pigmentation ranging from 1-10. We also classified colour phenotypes between 1-10, with *C. philodice* assumed to be 1-3, *C. eurytheme* ranging from 8-10 and intermediates represented by ranges 4-7. To standardize these categories and analyze the samples collected from fieldwork based on orange pigmentation, with intermediate *Colias* colours assumed to be hybrids, we developed a classification scale using museum samples from the Royal Ontario Museum. We examined approximately 2,500 samples of *C. philodice* and *C. eurytheme* and chose those samples representative of each stage in a colour gradient, where the most pale yellow individuals were classified as ‘1’ and the most orange were classified as ‘10’. Furthermore, we compared these representative butterflies to a wide range of paint chips ranging from light yellow to dark orange, so that each butterfly (1-10) could be represented as a paint colour. For fieldwork, we created a sampling sheet using these colour swatches to ensure consistency among different observers and environments. The response variables for the beta regressions include a scale for 0-1 which represent the proportion orangeness based on the original scale (Fig. 1; Table S1).

**Figure 1:**
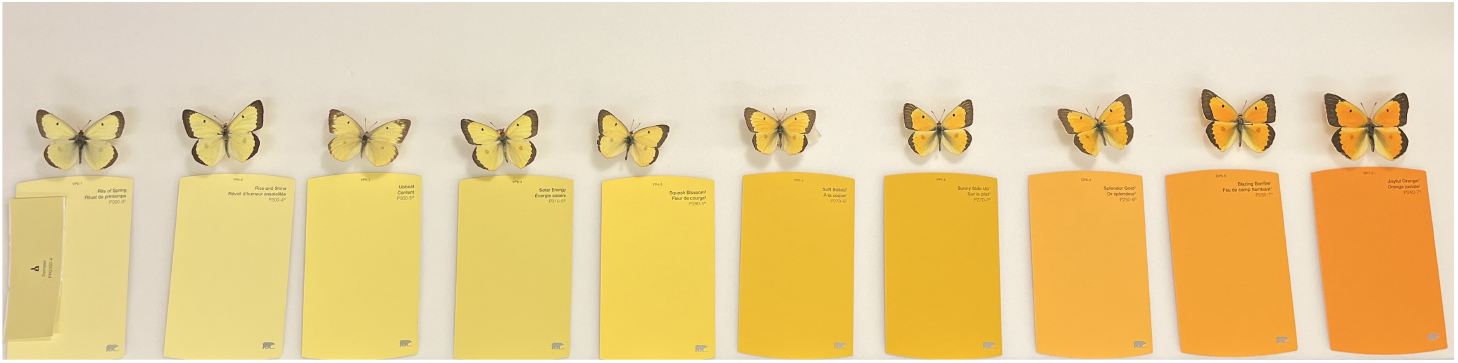
Objective numeric classification for *C. eurytheme* and *C. philodice* created using variable specimens from the Royal Ontario Museum. The colour swatches serve as a reference for the colour classification of all butterflies that were collected in the field. This standardized method allows for a framework to assess colour variation in *Colias*. Each *Colias* was then assigned a number from 1-10 based on the colouration on its upperside.

### Survey methods

We did all of our *Colias* sampling in Toronto, as urbanized areas in the city are expanding (Blais et al., 2000), and there is substantial habitat variation among parks within the city limits, and *Colias* are widespread (Fig. 2). To identify the city of Toronto parks that would best represent a wide range of urbanization, we examined all green spaces throughout the city (City of Toronto Parks, Forestry and Recreation 2024). The selected sites varied and included a wide range of urban habitats including urban parks, universities, forests, and meadows (Fig. 3). Using iNaturalist (https://www.inaturalist.org), habitats were determined based on 3 factors; if *C. philodice* had been present within the previous years, if *C. eurytheme* had been present within the previous years, and if there was vegetation that *Colias* preferred, such as alfalfa (iNaturalist 2024; Fig. 2). The region spans approximately 35.55 km north-south and 35.38 km east-west, demonstrating the large-scale coverage of the study area within the city.

**Figure 2:**
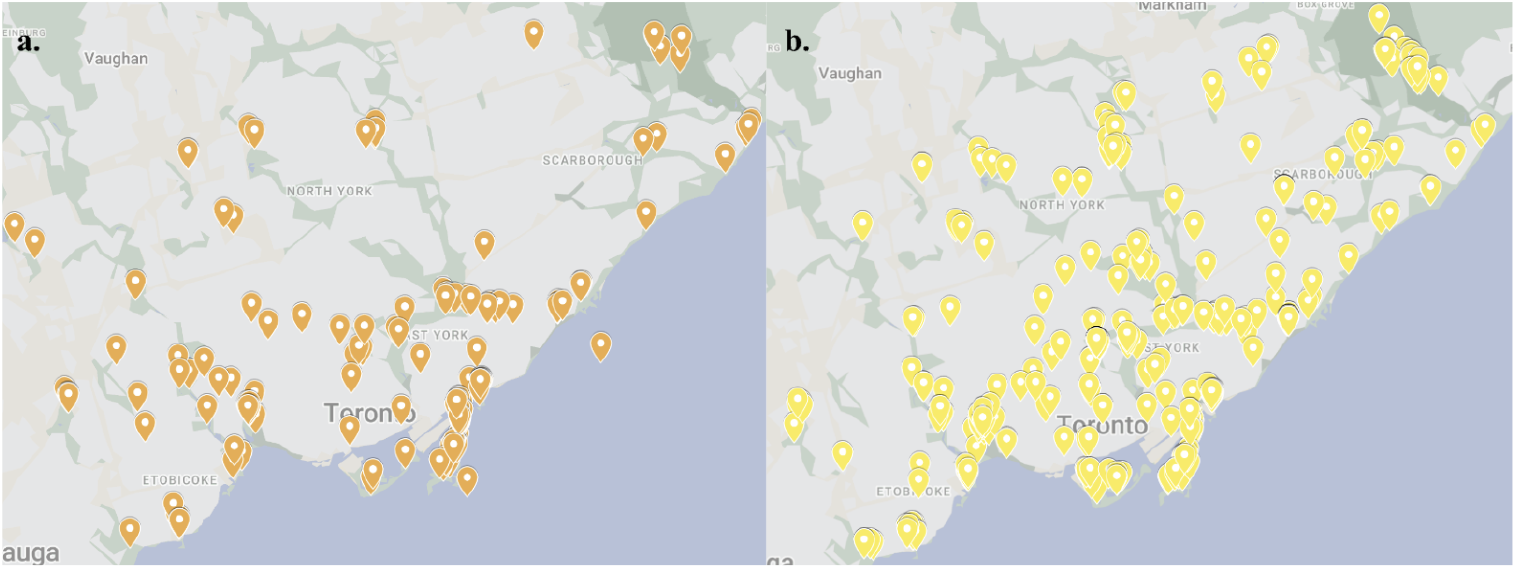
Comparing the historical observations (2012-2013) of *Colias* using iNaturalist. (a) Past *Colias eurytheme* observations in Toronto. (b) Past *Colias philodice* observations in Toronto.

**Figure 3:**
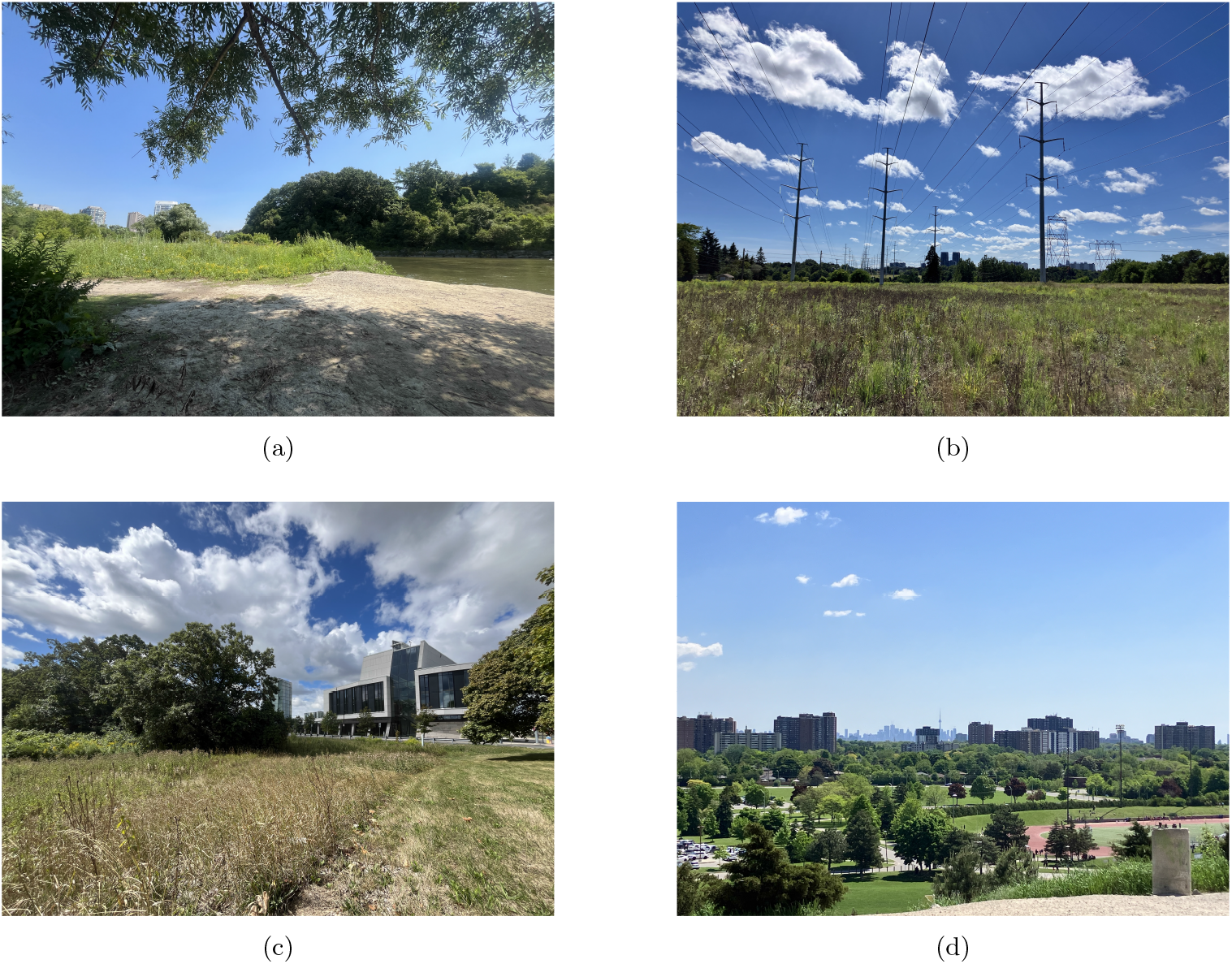
We selected sites to capture a mosaic of urban habitats. Shown here are some locations where we caught *Colias*: (a) the side of the Humber River with patches of meadow, (b) a hydro corridor featuring an overgrown meadow, (c) a university campus meadow, and (d) an urban park surrounded by buildings with the CN Tower in the background.

To systematically examine *Colias* in Toronto, we sampled between May 7, 2024, to July 23, 2024, haphazardly in City of Toronto parks and on university campuses. At each collection site, depending on the weather and visual inspection of the area, we sampled for between half an hour and three and a half hours. We attempted to catch all *Colias* seen using hand nets. For each butterfly that we collected, we identified them in the field using the sampling sheet, containing the standardized scale of colours created at the Royal Ontario Museum (Fig. 1). This allowed us to observe butterflies and phenotypically assign a value to each butterfly collected. We then inserted the collected butterflies into individual glassine envelopes and stored them in a −20 °C freezer for future genetic studies.

We assessed pedestrian density by counting the number of pedestrians within a radius of 3 metres, at each capture site, to calculate the index of human disturbance (Thaweepworadej and Evans, 2023). Furthermore, for each site, we estimated the distance to the nearest tree from the point where the butterflies were collected (meters). Julian date refers to the date of collection for each sample. To determine the various climate variables including temperature, we used the Government of Canada’s historical data reports (Environment and Climate Change Canada, 2024). This provided hourly updates using the Toronto Lester B. Pearson International Airport weather station (Environment and Climate Change Canada, 2024). We used estimates of ward population density (km^2^) and human population sizes from the City of Toronto (City of Toronto, 2025). Details of this are described below.

### Statistical analysis

All analyses were performed using `R v4.4.1` (R Core Team, 2024). We included a variety of data within our analysis as measures of urbanization. To determine the density of each Toronto ward, the city of Toronto ward profiles were used (City of Toronto, City Planning 2021). To create maps, we used the `R` packages `ggplot2, ggmap` and `sf` (Wickham, 2016; Kahle and Wickham, 2013; Pebesma, 2018; R Core Team, 2024). To determine the distances from each sampling point to the nearest road, we calculated the distance between butterfly capture locations and nearby roads in meters using the `R` packages `googleway, dplyr`, and `geosphere` (Cooley, 2023; Wickham et al., 2023; Hijmans, 2022; R Core Team, 2024).

To ask to what extent urbanization affects butterfly phenotypes we used the `R` package `glmmTMB` to fit a beta-regression model (Brooks et al., 2017; R Core Team, 2024). We used a beta-regression model as colour is being used as a proxy for hybrid score, which must always sum to one (100% ancestry) (Bolker, 2008). We scaled all variables and included distance to the nearest road, ward population density (km^2^), date of collection, 2021 ward population, distance to the nearest tree, pedestrians observed at the time of collection, and temperature. Additionally, parks were included as a random effect in the model to account for the expected variability between the different parks. Data visualization was facilitated by the `ggplot2` package (Wickham, 2016; R Core Team, 2024), which was used to generate scatter plots with regression lines for each predictor.

Finally, using the *ggpredict* function in R from the `ggeffects` package (Lüdecke, 2018; R Core Team, 2024), we generated model-based predictions across the full range of each scaled predictor variable. To isolate predictions to correspond to intermediate response probabilities, we applied custom filtering using the *dplyr* function (Wickham et al., 2023; R Core Team, 2024). This model extrapolates in which environments we would be most likely to find hybrids/intermediate phenotypes.

## Results

In total, we collected 122 butterflies from 12 sites (we visited 21; Fig. 4). We collected between 1 and 35 butterflies per site, and each location was visited at least twice (minimum number of visits =1, maximum =3). The average colour was 5.45 +/-1.70 (mode = 6, least common were 1 and 10) (Table 1).

**Table 1:** Different classification colours observed among *Colias*. Butterflies were ranked on a scale ranging from 1–10 (see Fig. 1). Individuals classified as 0 had the Alba phenotype, a white morph found exclusively in females with a Mendelian mode of inheritance (Tunström et al., 2023).

| Classification Colour | Hex Code | Number of *Colias* collected |
| --- | --- | --- |
| 0 | #FFFFFF | 2 |
| 1 | #FFEBA9 | 1 |
| 2 | #FFE99E | 2 |
| 3 | #FFE486 | 9 |
| 4 | #F7DA74 | 23 |
| 5 | #FFD467 | 17 |
| 6 | #FFB737 | 33 |
| 7 | #FAAB1C | 27 |
| 8 | #FFB14F | 4 |
| 9 | #FFA034 | 3 |
| 10 | #FA9335 | 1 |

**Figure 4:**
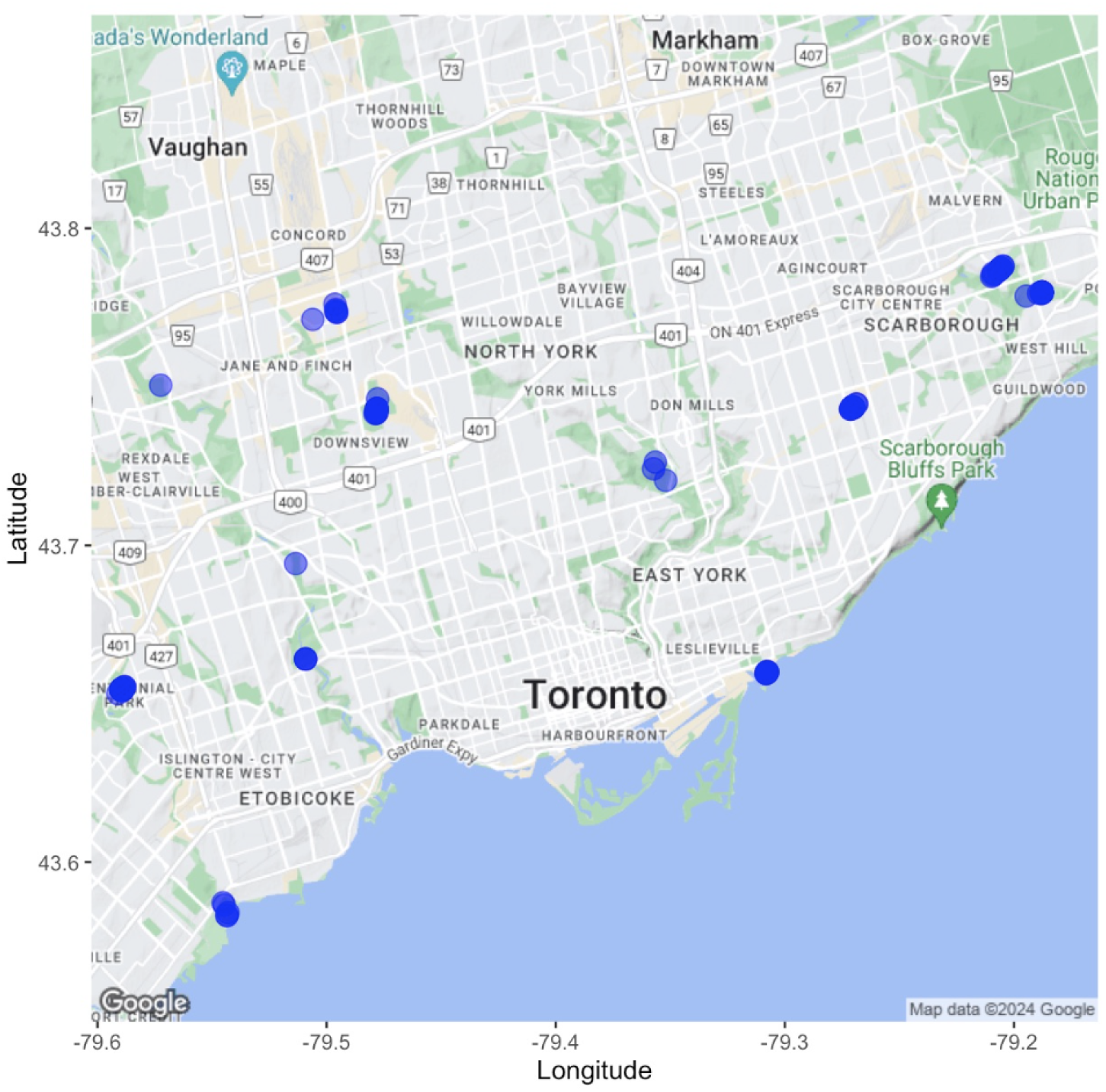
Map of Toronto displaying the amount and location of butterflies collected throughout the fieldwork period. Blue circles represent each *Colias philodice* and *C. eurytheme* that was collected.

Distance to road, Julian date, and pedestrians all were associated with the colour of butterflies in Toronto (Table 2). As distance to road increased, the proportion of orange butterflies observed also increased (Estimate = 0.267, z = 3.46, p *<* 0.001). Later in the season, the proportion of orange butterflies observed increased (Estimate = 0.263, z = 2.89, p = 0.004). Higher pedestrian activity resulted in a lower proportion of orange butterflies (Estimate = −0.383, z = −4.58, p *<* 0.001; Table 2). Other variables such as population density, 2021 population size, distance to nearest tree, and temperature, did not show statistically significant effects; however, temperature approached significance (Estimate = 0.165, z = 1.74, p = 0.082; Table 2, Fig. 5). The R ^2^ was 0.610 and the variance due to park was small (variance= 6.33 e-10). The variance of the random intercept for park number was 2.51 e-05.

**Table 2:** Results of the beta regression. Significant correlations are denoted with asterisks (*p *<*0.05; **p *<*0.01; ***p *<*0.001).

| Fixed effects | Estimate | Std. Error | z value | p(> \|z\|) |
| --- | --- | --- | --- | --- |
| Intercept | 0.0448 | 0.0712 | 0.629 | 0.529 |
| Distance to road | 0.267 | 0.0771 | 3.46 | 0.000541*** |
| Density population (km<sup>2</sup>) | 0.110 | 0.0881 | 1.25 | 0.213 |
| Julian date | 0.263 | 0.0910 | 2.89 | 0.003800** |
| 2021 population | -0.0817 | 0.0970 | -0.842 | 0.400 |
| Distance to nearest tree (m) | -0.0554 | 0.0769 | -0.721 | 0.471 |
| Pedestrians observed | -0.383 | 0.0835 | -4.58 | 4.5e-06*** |
| Temperature | 0.165 | 0.0949 | 1.74 | 0.0819 |

**Figure 5:**
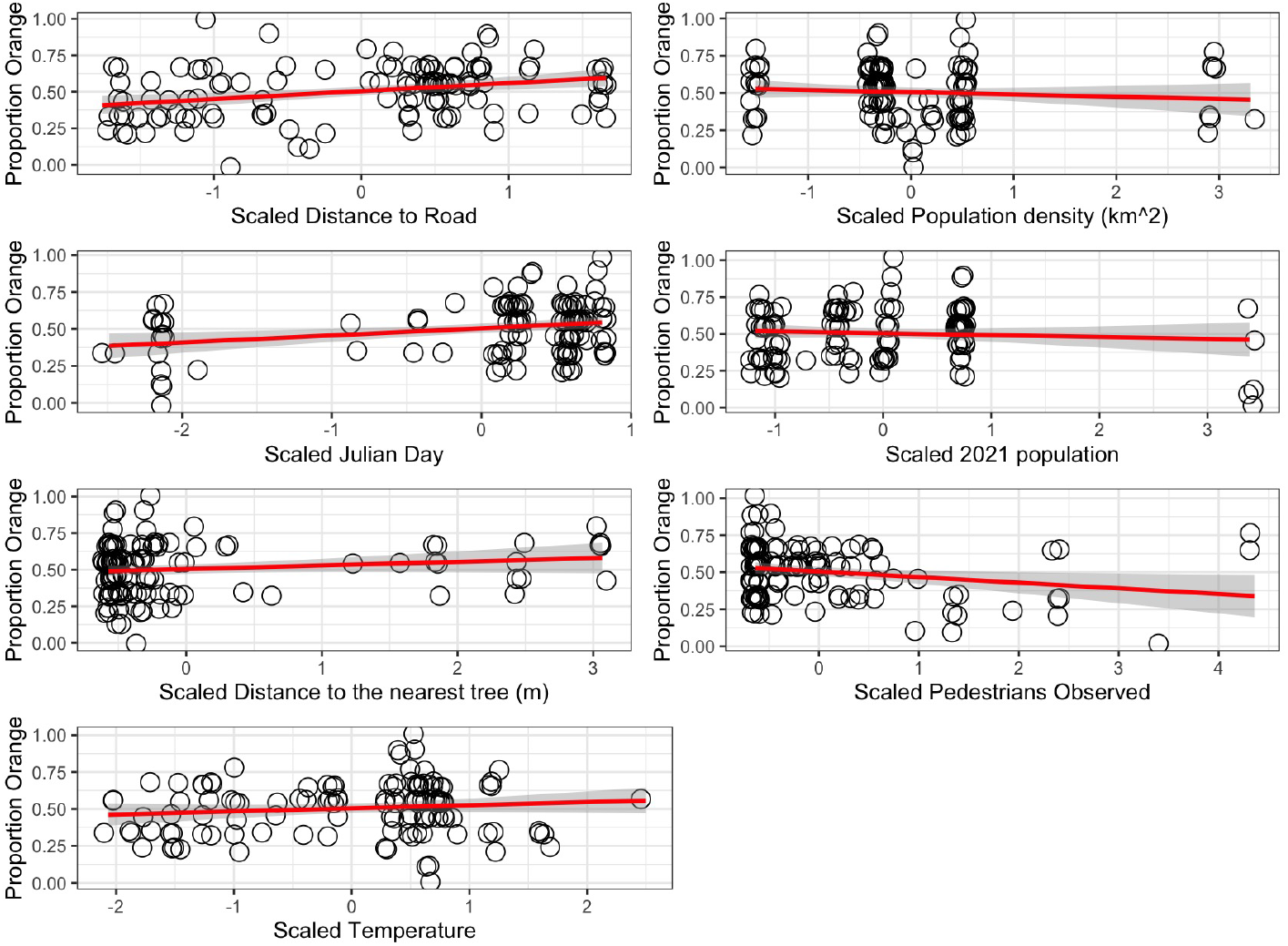
Relationship between the proportion of orange butterflies and distance to the nearest road, ward population density (km^2^), Julian days, 2021 ward population, distance to the nearest tree (m), pedestrians observed, and temperature. Proportion orange represents the butterfly colouration on a scale of 0-1 which represents the proportion orangeness based on the original scale (see Fig. 1). All variables on the x-axis are scaled. Each point represents a butterfly.

Distance to road, Julian date, and number of pedestrians appear to influence the probability of detecting hybrids. Using these model results, we then extrapolated predicted probabilities of hybrid occurrence across the observed ranges of the significant metrics. For example we might see a clouded-like hybrid (colour = 4, Fig. 1) in areas such as 8.08m away from the road, with 9.99 pedestrians, and on a julian day of 150 (May 29 of a leap year). We might see an orange-like hybrid (colour = 6) 350m away from the road, 0 pedestrians (which was rounded from −0.0193), and on a julian day of 205 (July 23 of a leap year).

## Discussion

We found that specific environmental and anthropogenic factors are correlated with the colour of *C. philodice* and *C. eurytheme*. We also developed a standardized colour scale for *C. philodice* and *C. eurytheme* identification. This new tool will allow for other studies to be consistent with ours when determining *Colias* colouration, enabling more uniform comparisons of phenotypic classification and consistency across studies.

We found that some of the urbanization metrics, including distance to road, Julian date, and number of pedestrians explain variation in *Colias* colour (Table 2). The distance to the nearest road has a significant effect on the proportion of orange *Colias* observed (Table 2). As the distance to the nearest road increases, the proportion of orange phenotypes increases, suggesting that *C. eurytheme* and their backcrosses are more likely to be observed farther from human disturbances such as roads. Sampling date had a significant effect on the proportion of orange *Colias* butterflies (Table 2), which is not surprising, as previous studies have demonstrated that colour deepens throughout the season(Gerould, 1943; Fenner et al., 2022). Pedestrian density at the collection sites showed a significant negative correlation with the proportion of orange butterflies (Table 2). This suggests that increased pedestrian activity may negatively impact the number of orange butterflies found in an area. The metrics of urbanization might affect *C. eurytheme* more than *C. philodice*, although work understanding the specific underlying mechanisms is needed. For example, it is unclear if there would be physiological differences between the species that would vary in response to sodium or lead, as has been shown to drive responses in other butterflies (Kemmerling et al., 2025; Snell-Rood et al., 2014). Future work could determine how the different butterflies are responding to the differences in environment.

We demonstrate here that *Colias* species can survive within urban habitats of varying levels of urbanization; however, variation occurs with urbanization, meaning that they are not randomly distributed. Other studies have found that urban areas are important for species and that the way species react varies. There are a number of species that are at higher abundance in cities when compared to more rural settings, including European Red Squirrel (*Sciurus vulgaris*; Jokimäki et al., 2017), a variety of bird species (e.g. house sparrow, *Passer domesticus*; European starling, *Sturnus vulgaris*; and rock doves, *Columbea livia*; Savard et al., 2000; Batten, 1972). *C. philodice* might be such a species, as we were more likely to find *C. philodice* closer to roads and when there were more pedestrians. They might not be negatively affected by human disturbance, at least relative to *C. eurytheme*. In constrast, there is a decrease in bird diversity as urbanization increases, as many varied habitats are eliminated (Savard et al., 2000; Batten, 1972). Pollinators are also faced with the increasing effects of urbanization, which tends to negatively affect the abundance of their populations (Liang et al., 2023). This demonstrates a need to study the impacts of urbanization on a variety of species as it’s difficult to make general predictions about overall outcomes. Assessing how urbanization affects not only abundance, but how it might interact with hybridization to affect rates of biodiversity demonstrates how complicated these questions are. Looking at landscapes of urbanization are important study sites to view the outcomes of hybridization as they can inform us of how anthropogenic hybridization affects various species (Allendorf et al., 2001).

### Possible caveats and future directions

Future studies on *Colias* should use genomics to assess colour phenotypes as a proxy for hybridization. Other studies have found that phenotypic approaches can be useful to start examining hybridization; however, incorporating a genomic perspective is essential, as it allows for further details regarding anthropogenic hybridization (McFarlane and Pemberton, 2019). Similar studies that have been conducted in various regions in Northern California and Nevada have used 363 samples of *C. eurytheme, C. eriphyle* and phenotypic intermediates for genetic analyzes (Dwyer et al., 2015). Jahner et al. (2025) found that, while most genetically identified hybrids between *C. eurytheme* and *C. eriphyle* were phenotypically intermediate, there were many phenotypically intermediate individuals that were not hybrids, suggesting substantial colour polymorphism in each species. Future work on how urbanization affects hybridization should assess how well colour works as a proxy for hybridization between *C. eurytheme* and *C. philodice*, as we have likely overestimated the number of hybrid individuals in our study.

While our study took into account several environmental variables, there are other factors that could contribute to environmental variation. In particular, *Colias* nectar use is linked to the composition of the surrounding flora, highlighting the importance of vegetation surveys to understand resource selection (Watt et al., 1974). Urban parks in Toronto have varying vegetation which could affect microhabitats and *Colias* resource selection. It is yet unknown if or how hybrid *Colias* are different from parental type *Colias* in their resource use, but this would be an interesting avenue for future research.

## Conclusion

Significant gaps remain in our understanding the impact of urbanization on butterfly populations, including with regards to *Colias*. Urbanization is one of the major drivers of anthropogenic environmental change, and studies exploring its effects on biodiversity are essential (Seto et al., 2012; Liang et al., 2023). This study investigates how urbanization and human-induced disturbances influence hybridization using intermediate colour as a phenotypic measure of *Colias philodice* and *C. eurytheme*. “Clouded-like” intermediate individuals are more tend to live near roads early in the season, while “orange-like” intermediate individuals are more likely farther away from roads, later in the season and with fewer pedestrians. By examining the role of environmental heterogeneity and habitat fragmentation, we have begun to understand the changing dynamics of species interactions in an urbanized landscape.

## Author Contributions

**Amanda Sabatino:** Conceptualization, Data curation, Formal analysis, Investigation, Methodology, Visualization, Writing – original draft, Writing – review & editing. **Joshua P. Jahner:** Conceptualization, Methodology, Resources, Writing – review & editing. **S. Eryn McFarlane:** Conceptualization, Funding acquisition, Methodology, Resources, Supervision, Writing – review & editing.

## Acknowledgments

We thank Yiwei Xu for her assistance in the field and Amanda Meuser for her help with the colour scale. We are grateful to Robert Tsushima, Gordon Fitch, and Jenna LeBlanc for feedback. We are thankful to Brad Hubley and the Royal Ontario Museum for the use of their *Colias* collection and Patricia Landry from Toronto Parks. We also thank Stompy the Bear for being the official theme song of this research project. This manuscript is the result of AS’s undergraduate thesis project at York University.

